# High-resolution omics and functional profiling of fatty-acid-driven metabolic modulation reveal enhanced physiological relevance of engineered heart tissues

**DOI:** 10.64898/2026.11.23.768630

**Authors:** Carla Cofiño-Fabres, Chloë E. Feijer-Kaal, Federica Conte, José M. Rivera-Arbeláez, Edgar E. Nollet, Jolanda van der Velden, Arno van Rooij, Hans J.C.T. Wessels, Verena Schwach, Marcelo C. Ribeiro, Robert Passier

## Abstract

**Background:** Human pluripotent stem cell-derived cardiomyocytes (hPSC-CMs) hold vast potential for cardiovascular disease modeling and drug discovery, yet their immaturity limits their ability to recapitulate adult cardiac physiology. Three-dimensional engineered heart tissues (3D-EHTs) provide a functional and more physiologically-relevant model for disease modeling and drug screening. However, EHTs typically retain an immature metabolic phenotype. To further enhance maturation, we investigated how fatty acid (FA) supplementation and physiological insulin concentrations influence metabolic maturation and its consequences for cardiac function, pharmacological responses, and hypoxic injury.

**Methods:** The effects of FA and insulin supplementation on 3D-EHT maturation were investigated using an integrated molecular and functional approach. Four medium variations were evaluated through functional tests, calcium handling assays, immunofluorescence, ATP quantification, gene expression analysis, high-resolution proteomics, and metabolomics to relate metabolic remodeling to tissue function and responses to pharmacological and hypoxic challenges.

**Results:** FA supplementation enhanced the metabolic maturation of EHTs, surpassing the maturation achieved with an already highly specialized cardiomyocyte medium. The combination of FA with physiological insulin concentrations promoted a shift toward oxidative metabolism, supported by metabolic profiling and associated molecular changes identified through proteomic analysis. This metabolic remodeling was accompanied by changes in contractile performance and calcium handling, altered responses to clinically relevant pharmacological agents, and increased susceptibility to hypoxic injury. Notably, drug concentration-response relationships shifted toward clinically relevant exposure ranges for several compounds. Together, these findings establish a link between metabolic maturation, molecular remodeling, and functional responses of 3D cardiac tissues to pharmacological and pathological challenges.

**Conclusions:** Our findings demonstrate that metabolic culture conditions influence not only the maturation state of hPSC-CMs in 3D-EHTs but also their responses to drugs and hypoxic stress. By integrating multi-omics profiling with functional and disease-relevant assessments, this study provides a framework for improving the physiological relevance of engineered cardiac tissues for disease modeling and preclinical drug evaluation.

## 1. INTRODUCTION

Human pluripotent stem cell (hPSC)-derived cardiomyocytes (hPSC-CMs) represent a virtually unlimited cell source for the generation of *in vitro* models to study cardiovascular diseases and expedite *in vitro* drug screening ^1^. However, hPSC-CMs exhibit a relative fetal rather than an adult cardiomyocyte (CM) phenotype, limiting their potential to accurately model complex adult cardiac (patho-)physiology and predict drug safety and efficacy. In the postnatal heart, an important metabolic transition occurs, characterized by a reduction in circulating glucose and lactate concentrations, a decrease in insulin levels and an increase in fatty acid (FA) concentration ^2,3^. Concurrently, the heart switches from a glycolytic to an oxidative metabolism that remains until adulthood, with FAs becoming the main energy source (60-70%), instead of glucose (20%) and lactate (10%) ^4^.

Given the critical role of this metabolic transition in CM maturation ^2,3^, several studies have investigated the effect of FA supplementation in culture media on the maturation of hPSC-CMs, as these cells predominantly rely on glucose as their main energy source. These studies have reported an enhancement of maturation traits promoted by FA in the medium, such as reduced cell circularity, enhanced sarcomere alignment, increased twitch force, improved calcium handling, increased mitochondrial content and oxidative capacity ^5–9^. However, many of these studies include additional variations in culture conditions, such as three-dimensional (3D) culture or the addition of biochemical cues, alongside FA supplementation. These variations can confound the true effect of FA on maturation, making it challenging to isolate the specific contribution of FA. Moreover, comprehensive analyses of the metabolic and functional properties of FA-matured hPSC-CMs in 3D culture systems, such as Engineered Heart Tissues (EHTs) ^10–12^, are often lacking. This is mainly due to the challenges in obtaining reliable readouts, such as spatial heterogeneity and technical limitations in tissue handling that compromise assay accuracy. Consequently, most available metabolic data are derived from two-dimensional (2D) formats or provide only partial insight into the effects of FA supplementation in 3D models.

Recently, studies using hPSC-CMs in 2D cultures under hypoxic conditions have shown their potential for modeling ischemia. However, the strong reliance of hPSC-CMs on anaerobic glucose metabolism renders them more resistant to hypoxia compared to adult CMs ^13^, leading to reduced accuracy in modeling ischemic events *in vitro*. Notably, it has been shown that FA supplementation without glucose in hPSC-CMs in 2D cultures rendered them more susceptible to cellular damage post-hypoxia compared to glucose-based hPSC-CM cultures ^14,15^. Additionally, 3D culture by itself improves biological complexity and maturation, enhancing the responsiveness of hiPSC-CMs to hypoxia, highlighting the need for more mature systems to model this cardiac event ^16^. Despite this significant progress, studies employing hPSC-CM 3D models that incorporate the metabolic switch from glycolytic to oxidative metabolism to model ischemic events remain scarce ^17^.

The ability to achieve a more mature state is not only critical for recreating oxidative-demanding insults but also for improving the prediction of drug efficacy. Accordingly, enhancing hPSC-CM maturity by culturing in 3D has proven essential for enhancing their predictive capability compared to 2D culture. Moreover, human-derived EHTs offer a promising platform for recapitulating key aspects of human disease ^6^, as well as predicting drug safety and efficacy. Increasing maturity of current hPSC-derived *in vitro* models is expected to further enhance their predictive power and translational relevance across applications ^18,19^.

Here, we present a comprehensive analysis of hPSC-CMs following maturation in the presence of FAs in a 3D-EHT format, providing a comparison between single-variable conditions at metabolomic, transcriptomic, proteomic and functional levels. We demonstrate that under appropriate conditions, FA supplementation promotes the maturation of hPSC-CMs by increasing oxidative metabolism, with insulin levels playing a significant role in modulating this metabolic switch. Moreover, we showcase the applicability of the model for studying ischemic injury and assess how drug response of metabolically matured EHTs recapitulates clinically relevant readouts. This crucial advancement represents a significant stride towards the establishment of advanced and standardized human cardiac models that more closely resemble the adult human heart. Such models will enable mechanistic investigation of cardiac diseases, improve clinically relevant *in vitro* drug screening, thereby expediting discoveries in cardiovascular research.

## 2. METHODS

### 2.1. Human pluripotent stem cell (hPSC) culture

The experiments were performed using the human embryonic stem cell (hESC) lines NKX2.5^EGFP/+^-COUP-TFII^mCherry/+^ (cardiac dual fluorescent reporter line, obtained from Schwach at al.^20^ and the human induced pluripotent stem cell (hiPSC) line WTC-11 (GM25256, Wild Type C, Gladstone Institute of Cardiovascular Disease, University of California, San Francisco, origin: 30-year-old Japanese male). All hPSC lines were maintained as undifferentiated colonies in Gibco Essential 8 medium (Thermo Fisher, A1517001) on vitronectin (Thermo Fisher, A31804)-coated 6-well plates (Greiner Bio-One, 657160). The hiPSC line was passaged with 0.5% RevitaCell supplement (Thermo Fisher, A2644501). All lines were cultured under aseptic conditions and were tested negative for mycoplasma. The master cell bank passage number of the COUP-TFII line was p16 and was kept in culture to p35. The master cell bank passage number of WTC-11 line was p40+7 and was kept in culture to p40+55.

### 2.2. Differentiation of hPSCs into CMs

Differentiation of hPSCs to CMs was performed as previously described ^21^. Briefly, one day before starting the differentiation, hPSCs were seeded at a density of 20-25×10^3^ cells per cm^2^ on 6-well plates coated with Matrigel (83 μg protein/mL) (Corning, 354230) in Essential 8 (E8) medium. After 24 h (day 0), mesodermal differentiation was induced by addition of Activin-A (20 - 30 ng/mL, Miltenyi 130–115-010), BMP4 (20 - 30 ng/mL, R&D systems 314-BP/CF) and WNT activator CHIR99021 (1.5 μmol/L, Axon Medchem 1386) in BPEL medium ^22^. At day 3, BPEL containing WNT inhibitor XAV939 (5 μmol/L, R&D Systems 3748) was used to refresh cells. On days 7 and 10, cells were refreshed with plain BPEL medium. At day 13, beating CMs were metabolically selected with a 4-day lactate purification step. Lactate purification medium consisted of previously described cardiomyocyte medium (CM-medium) ^21^ without glucose and with an additional 5 mM sodium DL-lactate solution (60%, Sigma Aldrich, L4263). At day 17, purified CMs recovered for three more days in CM medium consisting of the above-described lactate purification medium supplemented with 4.5 mM D(+)-Glucose (Millipore, 1083371000). Afterwards, cells were dissociated with TrypLE 10X (ThermoFisher, A1217702) and cryopreserved in freezing medium comprising 50% Knockout serum replacement (KOSR) (ThermoFisher, 10828028), 40% CM medium, 10% DMSO (Sigma-Aldrich, D2650) and 0.5% RevitaCell. Cell populations containing at least 90% of green fluorescent protein (GFP)-positive cells (representing NKX2.5-positive cardiomyocytes in hESC lines) or cardiac troponin T (for WTC-derived CMs) were used for downstream applications (Figure S1a-b). Cultured cells were maintained at 37°C, 21% O_2_ and 5% CO_2_ (when cultured in E8 and BPEL-based media) or 10% CO_2_ (when cultured in CM-medium).

### 2.3. Cardiac fibroblast expansion

Human adult cardiac fibroblasts (FBs) were purchased from Promocell (C-12375) and expanded according to their protocol ^23^. Briefly, a T175 cell culture flask (Greiner) was incubated (at 37°C and 5% CO_2_) with 12 mL of FGM-3 (Promocell, C-23130) for 30 minutes. After thawing at 37°C, the cells were transferred from the cryovial to a cell culture flask containing the FGM-3 with an additional 18 mL of FGM-3. The cells were passaged after reaching 70-90% confluence. This process was repeated until passage 11 was reached. Then, FBs were frozen at a final concentration of 150×10^3^ cells/0.5 mL in freezing medium. The freezing medium consisted of 50% KOSR, 40% FGM-3, 10% DMSO and 0.5% RevitaCell supplement.

### 2.4. Engineered heart tissue generation

The EHTs were generated as previously described ^11^, using the commercially available EHT platform provided by River Biomedics ^24^. In brief, to generate 3 EHTs per well, CMs from the NKX2.5-COUP-TFII or WTC line and FBs were thawed, mixed in a ratio of 100:3, and resuspended to a final concentration of 16.8 × 10^6^ cells/mL and mixed with an extracellular matrix consisting of fibrinogen, Matrigel, aprotinin and thrombin. The resultant EHTs contained approximately 2.5 × 10^5^ cells each. After fibrinogen-thrombin crosslinking, the corresponding experimental culture media were added. The specific medium was refreshed after 24h and, thereafter, every 2-3 days. We tested four versions of the culture medium. First, as control we used the previously characterized and highly specialized CM medium (named *CT High*) containing supraphysiological insulin concentration: DMEM (Sigma, St. Louis, MO, USA, D5030), 4.5 mM glucose, 5 mM DL-lactate solution, 0.5 mM sodium pyruvate, 0.19 mM sodium hydroxybutyrate, 0.5 mM L-carnitine, 1 mM creatine, 5 mM taurine, phenol red (0.011 g/L), 1X Trace elements (A, B, and C; Corning), 1X chemically defined lipids (Life Technologies, Waltham, MA, USA), 2 mM Glutamax, 0.45 mM α-thioglycerol, 0.01X ITS-X, 50 µg/mL ascorbic acid-2P, 0.5% Pen-Strep, 3.5 g/L sodium bicarbonate, 100 nM T3, 100 ng/mL Long R3 IGF-1, and 1 µM dexamethasone. Second, we used a variation of the CT High medium containing physiological (lower) insulin concentration (named *CT Low*). Specifically, CT Low medium formulation contains lower ITS-X (0.00005X), and is supplemented with 0.85 nM sodium selenite (Sigma-Aldrich, 214485), 0.68 nM transferrin (Sigma-Aldrich, T8158) and 0.33 µM ethanolamine (Sigma-Aldrich, E0135). Third, we tested a FA-supplemented medium with supraphysiological insulin concentration (named *FA High*). FA High medium consisted of the same CT High medium supplemented with 1.5% AlbuMax II Lipid-Rich BSA (Thermo Scientific, 11021037), 0.82 µM Biotin (Sigma-Aldrich, B4639), 5 µg/mL vitamin B12 (Sigma-Aldrich, V6629), 2 mM L-carnitine, and 5 mM creatine. Lastly, we tested a variation of FA High but containing physiological insulin concentration (named *FA Low*), containing lower ITS-X (0.00005X) and supplemented with 0.85 nM sodium selenite, 0.68 nM transferrin and 0.33 µM ethanolamine. EHTs were kept at 37°C, 21% O_2_ and 10% CO_2_.

### 2.5. Force measurements

Force of contraction and kinetics were measured between day 18 and 21 after tissue casting. Briefly, brightfield movies were recorded using a Nikon ECLIPSE Ti2 inverted microscope (RRID:SCR_021068), under controlled temperature and humidity (37°C, 10% CO_2_), using a high-speed camera at 100 frames per second (fps) with 2X magnification. Two platinum electrodes (Advent Research Materials) were used to electrically stimulate the EHTs using a custom-made pacing device at 1.5 Hz (10 ms biphasic pulses, 4-5 V/cm) for 20 seconds.

To assess drug responses, the initial contraction force of the tissues was measured under electrically paced conditions (2Hz, 10 ms biphasic pulses, 4-5 V/cm for 20 seconds). To evaluate positive inotropic response, isoproterenol (Sigma, I5627) was administered at 0, 0.001, 0.01, 0.1 and 1 µM, digoxin (Sigma, D6003) at 0, 0.01, 0.1, 0.3, 1, 3 and 10 µM, and dobutamine (Merck, D0676) at 0, 0.1, 0.3, 1, 3 and 10 µM. Negative inotropic effects were determined by administration of verapamil (Sigma, V4629) at 0, 0.001, 0.01, 0.1, 1 and 10 µM, flecainide (Merck, F6777) at 0, 0.1, 0.3, 1, 3 and 10 µM. Mavacamten MYK-461 (Selleck Chemicals S8861) was administered at 0, 0.01, 0.1, 0.3, 1 and 3 µM. Contractile force was measured and quantified for each EHT after 5 minutes of incubation with each concentration of drug. All drug-induced responses were measured under electrical stimulation at 2Hz (10 ms biphasic pulses, 4-5 V/cm for 20 seconds). Increase or reduction of contractile force was assessed with respect to control condition (0 µM) in each case.

Hypoxic experiments were carried out in a BioSpherix Xvivo system X3. Before the treatment, culture medium (CT High or FA) was incubated overnight at 0.1% O_2_ (hypoxia) or 21% O_2_ (normoxia) and 10% CO_2_. Next day, basal contraction forces were measured for each tissue, and subsequently EHTs were transferred to the pre-incubated medium and kept in hypoxic or normoxic conditions. After 10 hours, contraction force was immediately recorded. In hypoxic conditions, the plates were sealed with tape to prevent oxygen exposure.

Force kinetics analysis was performed by tracking and measuring the displacement of the pillars over time using a previously published, custom-made software ^25^.

### 2.6. Calcium kinetics

EHTs were loaded with 4 µM Fluo-8 AM (Abcam, AB142773) dye in HBSS buffer for 1 h at 37°C. Afterwards, tissues were washed once with pre-warmed HBSS and left in the corresponding medium for 30 minutes at 37°C prior to imaging. For recordings, EHTs were transferred to a customized imaging plate, while preserving them mounted on the pillars. EHTs were electrically stimulated at 1.5 Hz and imaged in a Nikon ECLIPSE Ti2 fluorescent microscope under temperature and humidity control (37°C and 10% CO_2_). Analysis was performed using a custom-made Python software.

### 2.7. Immunofluorescence analysis

Live/Dead staining was performed following the manufacturer’s recommendations (Thermo Fisher, L3224). Quantification was done using standard analysis in ImageJ software (v. 1.54p, Fiji). Whole-mount staining of EHTs was performed as previously described ^11^. Briefly, EHTs were fixed for 1h at RT in 4% paraformaldehyde (PFA), washed at room temperature with 0.3% Triton-× 100 (Sigma-Aldrich; 3 × 20 minutes), blocked for non-specific binding with 3% BSA (Sigma-Aldrich, A9418), 0.3% Triton-× 100 (Sigma-Aldrich, T8787), and 0.1% Tween20 (Merck, P9416) in PBS overnight at 4°C. The primary antibodies were added and incubated for 2 days at 4°C, consisting of: ACTN2 (1:800; Sigma-Aldrich, A7811), TOM20 (1:250; Abcam, ab186734), MYL2 (1:100; Sigma-Aldrich, HPA019763), cTnT (1:250; Thermo Fisher, MA512960) and CX-43 (1:250; Sigma-Aldrich, C6219). Then, the tissues were washed at room temperature with 0.3% Triton-× 100 (3 × 20 minutes) and secondary antibodies at a concentration of 1:500 (Alexa Fluor-488 (Thermo Fisher, A21441), Alexa Fluor-647 (Thermo Fisher, A31571), and DAPI were added overnight at 4°C, keeping the samples protected from light. The next day, the tissues were washed three times with PBS for 20 minutes each at room temperature. All incubations were performed on a shaker. Finally, EHTs were mounted on a microscope slide with a 0.25 mm spacer (SunJin Lab, IS216) for confocal imaging with a Zeiss LSM 880 microscope.

Calculation of mitochondrial intensity was performed on images captured by confocal microscopy of TOM20 and DAPI immunostaining, using standard analysis in ImageJ software (v. 1.54p, Fiji). Total intensity was normalized by the number of cells (DAPI-stained nuclei) per tissue. Five images were acquired for each EHT and averaged for downstream quantification.

### 2.8. Cellular ATP content quantification

The ATP content of the samples was measured using CellTiter-Glo® 3D Cell viability assay (Promega, G9681), according to the manufacturer’s recommendations. EHTs from each medium condition were transferred to a 96-well plate in 100 µL of their corresponding medium, either plain, containing oligomycin (oxidative phosphorylation inhibitor; 3.5 µM, O4876-5MG, Sigma-Aldrich) or 2-deoxyglucose (2-DG, glycolysis inhibitor; 100 mM, D3179, Sigma-Aldrich). The plate was incubated for 30 minutes at 37 °C and afterwards equilibrated to room temperature. Per well, 100 µL of CellTiter-Glo® 3D Reagent mixed with CellTox™ Green (Promega, G8741), according to the manufacturer’s recommendations, was added and mixed for 5-10 minutes on an orbital shaker to induce lysis. The plate was incubated at room temperature for 25 minutes to stabilize the luminescent signal, after which the luminescence and green fluorescence were recorded in a VICTOR3TM 1420 Multilabel counter plate reader. After subtraction of the blanks, the total ATP content, and the contribution of glycolysis and oxidative phosphorylation to ATP production could be determined and corrected by green fluorescence signal per tissue (an indicative of the amount of lysate).

### 2.9. Real-time quantitative PCR

EHTs were removed from the pillars using tweezers, transferred to sterile microtubes and snap frozen in liquid nitrogen. Total RNA was isolated from each individual EHT using TRIzol reagent (FisherScientific; 15-596-026), according to the manufacturer’s instructions. EHTs in TRIzol were homogenized with a 28-30 gauge needle and syringe. The resulting RNA samples underwent reverse transcription to cDNA using iScript cDNA Synthesis kit (Bio-Rad; 1708891). Gene expression was assessed using a Bio-Rad CFX384 real-time system with SensiMix SYBR (Meridian Bioscience; QT615-05). The samples were analyzed with Bio-Rad CFX Manager software and normalized to the housekeeping human gene *RPLP0*. Primer sequences used are listed in Table 1.

**Table 1.**
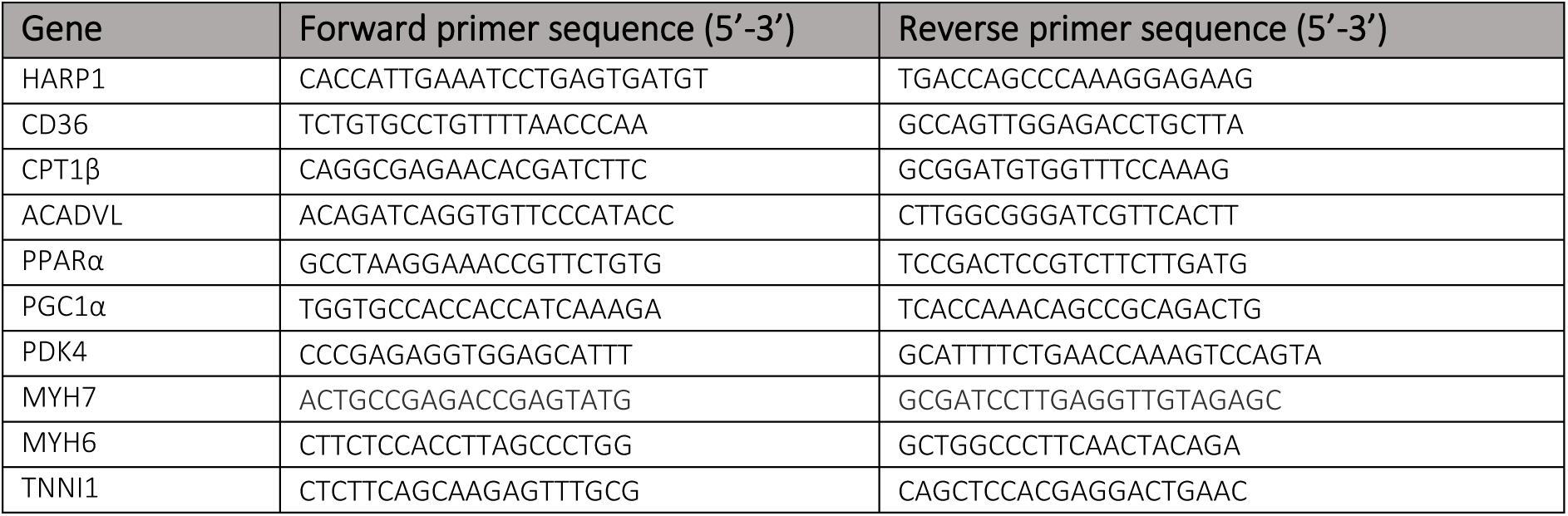
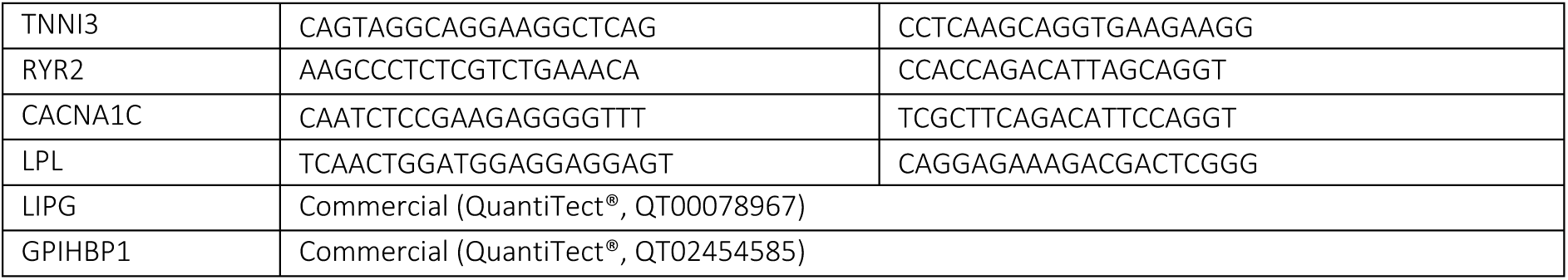
Primer sequences.

### 2.10. Mass spectrometry-based metabolomics

After force of contraction measurement, EHTs were washed twice in washing buffer (75 mM ammonium carbonate in water, with pH adjusted to pH 7.3±0.1 with acetic acid) and then incubated in extraction buffer (40:40:20 Methanol:Acetonitrile:water) for 10 minutes, as described in ^26^. After incubation, the tissue was removed and the extraction buffer containing polar metabolites was collected in a 1.5 mL Eppendorf tube and centrifuged at 14,000 rpm (or higher) for 5 minutes at 4 ᵒC. The extracts were then dried overnight (15 hours) in a Savant SC100 RVT 100 vacuum centrifuge (equipped with an external oil pump) at room temperature. Afterwards, the dried metabolic extracts were stored at −80 °C until analysis. Upon analysis, the metabolic extracts were resuspended in 100 μL of MS-compatible ultra-pure water (VWR Chemicals, HPLC-super high gradient, 83650.320).

For the measurement of phosphate sugars, 25 μL of each extract were injected into an Agilent 1290 ultra-high performance liquid chromatography mass spectrometer (UHPLC) coupled with an Agilent 6490A tandem quadrupole mass spectrometer (UHPLC-MS/MS), run in multiple reaction monitoring (MRM) mode. Chromatographic separation was obtained using a 0.25 μL/minute flow rate and a gradient from 0 to 100% mobile phase B over a 25-minute total run time. Mobile phase A (12 mM acetic acid, 10 mM tetrabutylammonium, 2 mM acetylacetone, 3% methanol in water) and B (12 mM acetic acid, 10 mM tetrabutylammonium, 2 mM acetylacetone, 3% methanol, 80% acetonitrile in water) were used to separate eight phosphate sugars (Figure S3d). Other analysis settings and the acquisition method are described in ^27^.

For the measurement of TCA cycle metabolites, 20 μL of each extract were diluted with 20 μL of stable isotope solution, containing 3 µM U-^13^C_6_-citrate, 6 µM U-^13^C_5_ 2-ketoglutarate and 3 µM of U-^13^C_4_ succinate, and then centrifuged for 3 minutes at 2000 rpm at 15 ᵒC, and shaken for 3 minutes at 100 rpm. Then, 1 µL of each extract was injected into a Waters I-Class Acquity fitted with a peek-lined Inert Sustain AQ-C18 (2.1×100 mm, packed with 3 µM particles, 100Å pore size) column connected to a Waters Xevo TQS micro mass spectrometer. The TCA cycle metabolites were separated through a linear gradient of 10 minutes (plus 1 additional minute during sample injection) with a flowrate of 0.25 ml/min, according to the method previously described ^28^. The gradient was run as follows: 3.5 minutes (conditioning time), 1.5 minutes with 95% phase A and 5% phase B, 1.5 minutes with 75% A and 25% B, 3 minutes with 100% B, 2.5 minutes with 100% A.

The peaks of the chromatograms from both methods were integrated via Skyline (v.20.2) software, using the transitions lists described in ^29^. The resulting peak areas were normalized based on the sum of all included peaks (also known as ‘total peak area’ normalization). The normalized data were processed and statistically analyzed in GraphPad PRISM (v.9.4.1), as described in paragraph ‘Statistics’ below. After testing the normality of data distributions, we opted for Kruskal-Wallis test (without correction for multiple comparisons), with a p≤0.05, to evaluate the statistical significance of observed metabolite differences between sample groups.

### 2.11. Quantification of fatty acid oxidation

EHTs were removed from their pillars and placed in ice-cold preservation solution (BIOPS solution), containing: K2EGTA (7.2 mM), CaK2EGTA (2.8 mM), ATP (5.8 mM), MgCl_2_ (6.6 mM), taurine (20 mM), phosphocreatine (15 mM), imidazole (20 mM), dithiothreitol (0.5 mM) and 2-(N-morpholino)ethanesulfonic acid (MES; 50 mM). The pH of the solution was adjusted to 7.1 with potassium hydroxide (KOH). EHTs were permeabilized using saponin (50 µg/mL) in ice-cold preservation solution for 25 minutes and subsequently washed twice for 10 minutes in ice-cold mitochondrial respiration medium (MiR05), containing EGTA (0.5 mM), MgCl_2_ (3 mM), potassium lactobionate (60 mM), taurine (20 mM), KH_2_PO_4_ (10 mM) HEPES (20 mM), sucrose (110 mM) and 1 g/L fatty acid free bovine serum albumin. The pH of the solution was adjusted to 7.1 with KOH. Per treatment condition, 2-4 EHTs were pooled and transferred to a measurement chamber of a high-resolution respirometer (Oxygraph-2k; Oroboros Instruments, Innsbruck, Austria). To assess functional changes in fatty acid oxidation, we measured the capacity to oxidize the activated fatty acid octanoylcarnitine (C8:0), as described previously ^30^. First, a low concentration of malate (0.1 mM) and ADP (5 mM) were added. Malate is converted to oxaloacetate by malate dehydrogenase, which subsequently condenses with β-oxidation-derived acetyl-CoA into citrate, the latter needed to prevent accumulation of acetyl-CoA and concomitant inhibition of β-oxidation. Next, cytochrome-c (10 µM) and octanoylcarnitine (0.2 mM) were added. Baseline respiration induced by malate and ADP was subtracted from all values as background. All experimental protocols were carried out at 37 °C, with oxygen levels maintained above 300 µM throughout the experiment to avoid limitations in oxygen supply.

### 2.12. Mass spectrometry-based proteomics

After force of contraction measurement, 3D EHTs were washed in DPBS 1X (without calcium and magnesium), (Thermo Fisher, 14190144). Next, the tissues were removed from the plastic pillars using tweezers, collected into 1.5 mL Eppendorf microtubes and snap-frozen by submerging each microtube in liquid nitrogen. The tissues were then homogenized in 20 µL 8 M urea/10 mM Tris (pH 8.0) using disposable plastic pestles (Cytiva, 80648337). The homogenates were incubated with 20 µL of reducing buffer (10 mM dithiothreitol in water) for 30 minutes at room temperature, and later with 20 µL of alkylating buffer (50 mM 2-chloroacetamide in 50 mM ammonium bicarbonate) for 1.5 hours at room temperature in the dark. Next, 20 µL of ammonium bicarbonate and 2 µL of Trypsin (5 µL/20 µg protein, Promega, V511C) were added to the homogenates, followed by overnight incubation at 37 °C. After peptide quantification, each sample was diluted in MS-compatible ultra-pure water to ensure the loading of a protein amount of 100 ng in a 20 µL volume for each EVOTIP Pure tips (Evosep, EV2013), according to manufacturer’s instructions. Tryptic digests were analyzed by nanoflow liquid chromatography (Evosep One, Evosep Biosystems) coupled online to a trapped ion mobility spectrometry-quadrupole time-of-flight mass spectrometer (timsTOF Pro2, Bruker Daltonics) via a nanoflow electrospray ionization source (CaptiveSprayer, Bruker Daltonics). Peptide separations were performed via C18 reversed phase liquid chromatography (Evosep EV1137 30SPD performance column; 150 mm length x 0.150 mm internal diameter, 1.5 µm C18AQ particles) using the pre-programmed 30 samples per day (30SPD) Evosep One method. The mass spectrometer was operated in positive ionization mode using the default data independent acquisition-Parallel Accumulation SErial Fragmentation (dia-PASEF) method ^31^. The instrument method parameters were as follows: 100-1700 m/z mass range, 0.6-1.6 1/K0 mobility range, 32 mass steps per cycle, 100 msec ramp time (100% duty cycle), 26 Da mass width, 1 Da mass overlap cycle, 0 mobility overlap, 1 mobility window. Acquired spectra were processed in real-time by a PaSER box (Bruker Daltonics) using the Data Independent Acquisition-Neural Network workflow (DIA-NN v2023) ^32^ with the following parameters: database used: Uniprot (incl. isoforms) from H. sapiens, precursor FDR 0.01%, precursor mass tolerance 15 ppm, MS2 mass tolerance 15 ppm, protein inference: protein name (FASTA) and Top N3 label-free quantitation (LFQ). The spectral library used was pre-constructed from data dependent acquisition-Parallel Accumulation SErial Fragmentation (dda-PASEF) measurements ^33^ of 2D cultured myoblasts, 3D EHTs, and human heart biopsy material that were searched against the Uniprot canonical and isoform protein human sequence database using the PaSER ProLuCID workflow. ProLuCID and DTASelect settings: 20 ppm precursor ion mass tolerance, 30 ppm fragment ion mass tolerance, carbamidomethyl (Cys) as a fixed modification, N-terminal acetylation and methionine oxidation as variable modifications, strict tryptic cleavage allowing for 1 missed cleavage, 6 amino acids as minimum peptide length, and 1% protein-level false discovery rate cutoff.

### 2.13. Statistics

Statistical analysis and visualization were performed using GraphPad Prism 8, and in-house Python3 scripts based on Python libraries (including NumPy, Pandas and Matplotlib), run in Spyder IDE (v6.0.4). The specific tests used for assessing statistical significance in the different assays are indicated in the caption of each figure. Parametric tests (one-/two-way ANOVA, Student’s t test for paired or unpaired analysis, Welch’s t-test) and non-parametric tests (Kruskal-Wallis test) were applied accordingly to evaluate the differences in means between groups and/or conditions. Dose-response relationships were analysed by nonlinear regression using a four-parameter logistic equation (variable slope). Raw data were normalized prior to fitting: for agonists, responses were scaled between 0% (smallest mean) and 100% (largest mean); for inhibitors, scaling was performed inversely. EC₅₀ or IC₅₀ values were obtained from the fitted curves. Results are displayed as mean ± standard error of the mean (SEM) unless stated otherwise and were considered significantly different with p≤0.05.

## 3. RESULTS

### 3.1. Fatty acid-based medium reduces tissue thickness without cell death

Our objective was to investigate the effect of FA supplementation in our previously characterized and highly specialized CT High ^34^ using our previously described EHT platform, which is compatible with a commercial 12-well-plate format ^11^. For this study, flexible plastic pillars, produced by River BioMedics, were used instead of PDMS pillars to enable high-throughput fabrication via injection molding (Figure 1a) ^25^. hPSCs were differentiated to CMs using an established protocol ^21^, including a 4-day lactate purification step. The purified CMs were harvested and frozen at day 20. As previously described, EHTs were generated in a fibrinogen-thrombin matrix ^11^ containing purified CMs (Figure S1a-b) and 3% supportive FBs. Tissues were cultured for 2-3 weeks in control or FA-supplemented media (Figure 1b). Since insulin concentration (17 nmol/L) in the control medium exceeded physiological levels, potentially interfering with FA metabolism ^4^, we lowered the insulin concentration to 85 pmol/L and compared the effect in both control and FA media. This resulted in four experimental conditions: control and FA media, each with either high or low insulin levels (Figure 1b). The FA medium was based on the existing CT High medium formulation, which includes physiological calcium levels, low glucose (4.5 mM), lactate and other components fulfilling the energetic demands of cardiomyocytes (e.g. L-carnitine, taurine, creatinine). FA supplementation was achieved using AlbuMAX, a mixture of linoleic acid, oleic acid and palmitic acid bound to albumin. The concentration was based on the recommended manufacturer’s level (183 µM) ^35^, and as previously reported ^9^. However, EHTs exposed to this concentration showed no significant changes in contraction force or expression of metabolism-related genes, regardless of insulin levels (Figure S1c-f). Consequently, we hypothesized that increasing AlbuMAX concentration to approximate the FA levels found in adult plasma ^36^ may prompt the necessary metabolic switch. Therefore, we proceeded to increase the AlbuMAX concentration to 1.5% (w/v) (550 µM) in both FA High and FA Low groups for subsequent investigations.

**Figure 1.**
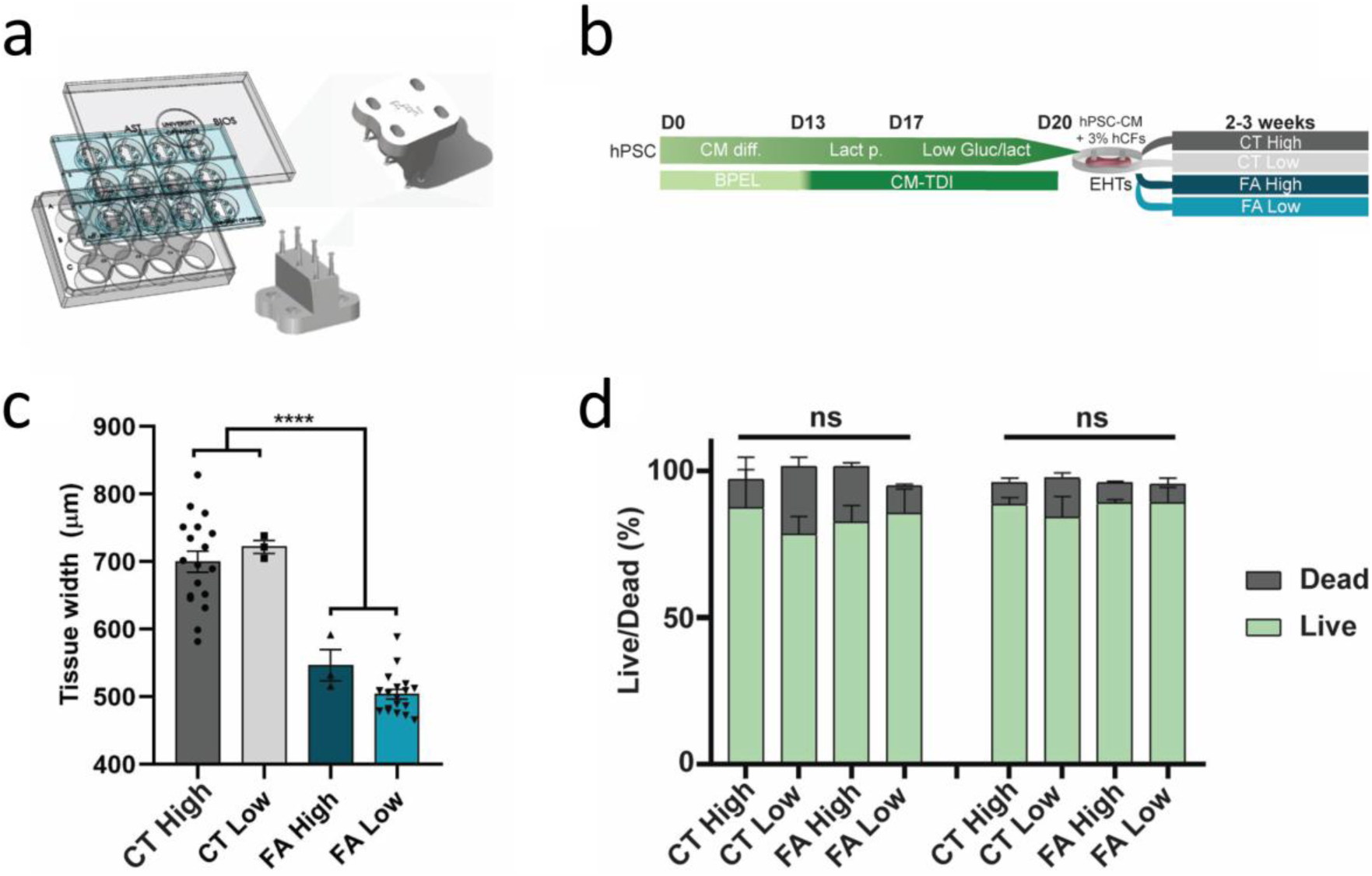
Fatty acid supplementation promotes a reduction in tissue width due to compaction without compromising tissue viability. (a) EHT platform used for the study, consisting of a modular 12-well plate platform design with flexible plastic pillars placed in one holder per well. (b) Experimental design illustrating the schematic differentiation of hPSCs towards cardiomyocytes (hPSC-CMs) in BPEL medium, following lactate purification and recovery in cardiomyocyte specific medium (CM-TDI) till freezing at day 20. During EHT casting, hPSC-CMs are mixed with 3% human cardiac fibroblasts (hCFs) and cultured for 2-3 weeks in four different media conditions (CT High, CT Low, FA High and FA Low). (c) Tissue width of EHTs after 20 days of culture in each medium condition (n=3-18 tissues from 3 CM batches). (d) Quantification of LiveDead images in each medium condition at day 5 and 20 after tissue formation (n = 3 tissues/timepoint from 3 CM batches). Abbreviations: CT High, control medium with high insulin; CT Low, control medium with low insulin; FA High, FA medium with high insulin; FA Low, FA medium with low insulin. (c-d) One-way ANOVA with post-hoc correction via Tukey test. Data presented as mean ± SEM. ****p<0.0001; ns = not significant.

We found that culture in FA media led to a reduction of tissue width (Figures 1c and S2a-b). To exclude that this decrease in width was associated with FA-induced toxicity due to the increased FA concentration, we evaluated the viability of the hPSC-CMs in the tissue after 5 and 20 days of medium conditioning (Figures 1d and S2c). Across all media conditions and time points, viability was comparable, ranging from 78% to 90%. These findings indicate that FA supplementation promotes tissue compaction without inducing cell loss.

### 3.2. Fatty acids and physiological levels of insulin induce a switch to oxidative metabolism

To evaluate the metabolic state and energy supply mechanisms of the EHTs, we quantified ATP production through different metabolic pathways. Since the energy (ATP) demands of CMs are coupled to their structural conformation (e.g. 2D vs 3D cultures), we assessed their metabolic activity in the 3D configuration, maintaining the physiological environment in which they are cultured. For that, we made use of a bioluminescent ATP assay (CellTiter-Glo®), which allowed us to preserve the EHTs on their flexible pillars until the moment of measurement, thereby sustaining their contractile and metabolically active state. Since the amount of ATP is proportional to the number of live cells per tissue ^37^, we corrected it by normalizing to the amount of DNA per EHT, with a DNA-binding dye (CellTox-Green Dye™). We validated the correlation between these two variables using increasing cell densities (Figure S3a-c). Next, we measured the ATP levels of the EHTs in each of the four media conditions, following treatment with either 2-DG (inhibitor of glycolysis) or oligomycin (inhibitor of oxidative phosphorylation (OxPhos) activity), and compared the resulting ATP levels with basal ATP levels measured without inhibitors (Figure 2a). As expected, EHTs cultured in control medium, either with high or low insulin levels, exhibited a marked reduction in ATP content following 2-DG treatment, but not with oligomycin, indicating that the hPSC-CMs predominantly rely on glycolysis for ATP synthesis. Interestingly, EHTs cultured in FA medium with high insulin did not show significant difference in glycolysis and OxPhos contribution to ATP generation, although a trend towards a greater contribution of glycolysis than OxPhos was observed. However, upon reduction of insulin in FA-supplemented medium to physiological levels, a reversed trend was observed. Specifically, EHTs treated with oligomycin exhibited a greater decrease in ATP levels compared to those treated with 2-DG, suggesting a metabolic shift from glycolysis to OxPhos (Figure 2a).

**Figure 2.**
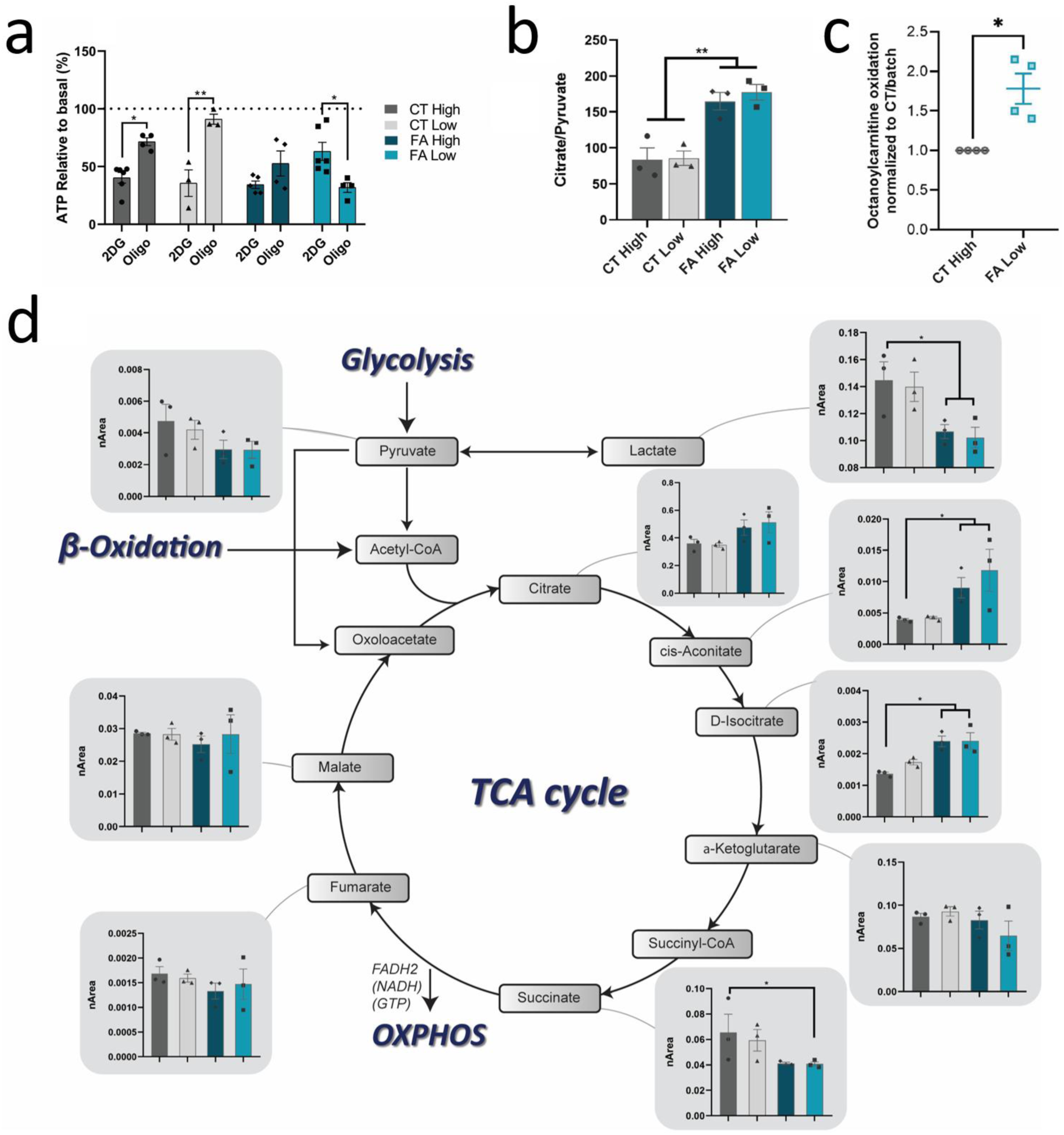
Metabolic analysis of EHTs reveals that FA supplementation with physiological insulin levels increases TCA activity and promotes FA β-oxidation over glycolysis. (a) Values of ATP after treatment with 2-Deoxyglucose (2-DG) or Oligomycin (Oligo) relative to the basal ATP per media condition (dashed line), normalized per tissue lysate. Two-way ANOVA with Tukey post-hoc test (n = 3-6 tissues from 3 CM batches). (b) Schematic representation of the glycolysis and tricarboxylic acid cycle (TCA) pathways and the corresponding bar graphs of the metabolites measured via liquid chromatography mass spectrometry (LC-MS/MS) in EHTs. Dark grey = CT High; Light grey = CT Low; Dark blue = FA High; Light blue = FA Low. Kruskal-Wallis test (n=3 tissues from 3 CM batches). (c) Ratio of TCA over glycolysis, determined by the ratio of citrate and pyruvate metabolites measured by mass spectrometry, respectively. One-way ANOVA with Tukey post-hoc test (n=3 tissues from 3 CM batches). (d) Fatty acid β-oxidation capacity measured by octanoylcarnitine oxidation, in CT High and FA Low treated EHTs, and normalized to CT High. Welch’s t-test (n=4 tissues from 4 CM batches). Data plotted as means ± SEM. *p<0.05, **p<0.01.

### 3.3. Metabolomic analysis reveals FA medium supports higher TCA activity and FA β-oxidation

To quantify the effects of FA and insulin treatment on the individual metabolic pathways for ATP generation, we performed mass spectrometry-based metabolomic profiling (Tables S1 and S2). To avoid disrupting the cell’s metabolism, we employed a recently established protocol to extract polar metabolites from intact EHTs with minimal handling, by extracting metabolites while tissues are kept on pillars ^26^. The resulting metabolic extracts were used for targeted metabolomics analysis to quantify hexose phosphates, sugar derivatives and metabolites from the tricarboxylic acid (TCA) cycle. In accordance with the ATP assay results, pyruvate and lactate, two products of glycolysis, were found in higher amounts in EHTs treated with control media compared to FA media (independent of insulin concentration) (Figure 2b). On the other hand, TCA cycle metabolites, such as citrate, cis-aconitate or D-isocitrate were higher in EHTs cultured in FA-supplemented media (Figure 2b). Decreased succinate levels in FA-treated EHTs may indicate enhanced OxPhos, as the reduction in succinate is often associated with increased mitochondrial activity, leading to accelerated oxidation of succinate to fumarate ^38^. Additionally, we used the ratio of citrate (first TCA metabolite derived from acetyl-CoA) to pyruvate (end product of glycolysis) as a readout for TCA cycle activity relative to glycolysis-derived lactate production, as previously reported ^39,40^. Notably, FA treatment (independent of insulin concentration) increased the citrate/pyruvate ratio, indicating enhanced TCA cycle activity (Figure 2c). Given that FA treatment with physiological insulin (FA Low) induced a metabolic shift towards a more oxidative metabolism and exhibited a modest increase in TCA cycle activity compared to FA with high insulin (FA High), we selected this condition for subsequent analyses and compared the control medium (CT High) with the best-performing condition (FA Low).

To further validate these findings, we performed a fatty acid β-oxidation assay on EHTs cultured in CT High or FA Low medium. This revealed a strong increase in β-oxidation capacity in the FA Low condition, supporting the hypothesis that increased TCA cycle activity was driven by enhanced β-oxidation (Figure 2d). Finally, we calculated the adenylate energy charge (AEC) to infer the EHT’s energy status under the two conditions ^41–43^. The AEC was comparable between conditions, indicating that FA Low medium does not impair overall cellular energy status (Figure S3e). Taken together, these findings suggest that FA Low medium promotes a more oxidative mode of energy metabolism to meet cellular energy demands.

### 3.4. EHTs in FA Low medium exhibit increased expression of genes and proteins involved in the TCA cycle, β-oxidation and oxidative phosphorylation

Metabolomic analysis revealed that FA supplementation with physiological levels of insulin enhanced the metabolism of hPSC-CMs, mainly through increased β-oxidation and OxPhos. To uncover molecular mechanisms underlying these changes, we analyzed the gene expression of metabolism-related genes and performed proteomics profiling (Table S3) on EHTs treated with FA Low versus control (CT High) (Figure 3). FA Low medium promoted the upregulation of genes involved in various steps of fatty acid metabolism, including FA uptake (*CD36*), mitochondrial transport (*CPT1B*) and FA β-oxidation (*ACADVL*), as well as transcription factors known to enhance FA catabolism (*PPARA*, *PGC1A*, *PDK4*) (Figures 3b and S4a-c). Comparison with FA High-treated EHTs further confirmed that the observed upregulation of these genes was triggered by the combination of FA addition and reduction of insulin concentration (Figure S4d).

**Figure 3.**
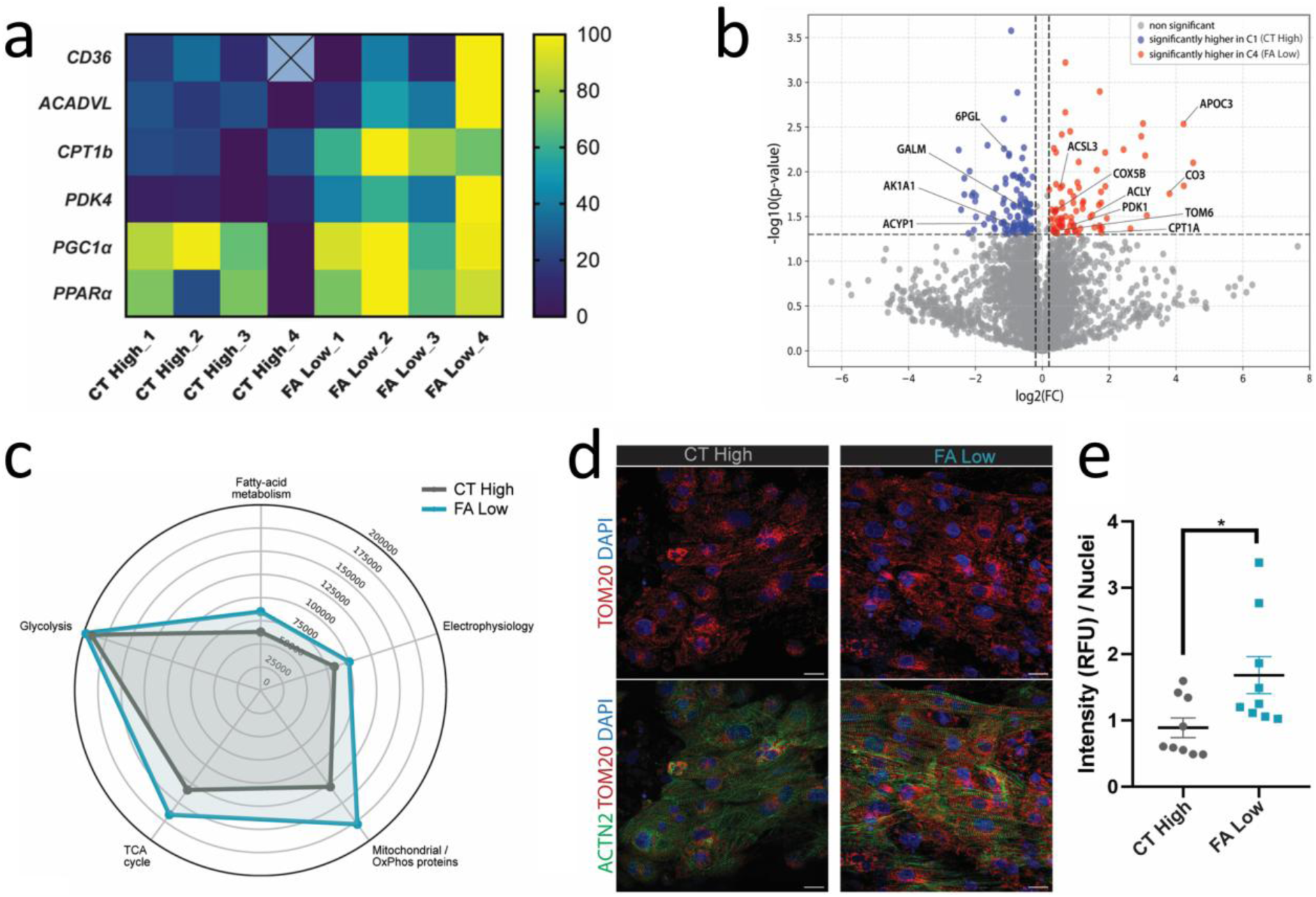
Gene expression and proteomic profiling indicate that FA supplementation and physiological insulin concentration enhance FA uptake, β-oxidation, TCA cycle and oxidative metabolism in EHTs. (a) Heatmap depicting the expression, measured by RT-qPCR, of genes involved in FA uptake and metabolism (n=4 tissues from 4 hiPSC-CM batches; cross indicates missing sample). (b) Volcano plot depicting differential protein expression as measured via LC-MS/MS (n = 3 tissues from 3 CM batches). The volcano plots were generated for exploratory analysis using the Welch’s T-test (p=0.05, t=0.2) applied to the normalized, log2-transformed proteomics dataset (∼4500 proteins/EHT). (c) Radar plot of biological processes across experimental groups based on average LFQ values of respective proteins. (d) Representative confocal images of the mitochondrial marker TOM20, alpha-actinin (ACTN2) and nuclei (DAPI) in CT High- and FA Low-treated EHTs. Scale bar = 20 µm. (e) Quantification of mitochondrial content based on TOM20 intensity normalized per nuclei per picture. Welch’s t-test (n=8 tissues, 5 images per tissue). Data plotted as means ± SEM. *p<0.05.

Exploratory proteomic profiling revealed the upregulation of proteins generally involved in lipid metabolism and mitochondrial energy production (such as CO3, CPT1A, APOC3, ACLY, COX5B, ACSL3, PDK1 and TOM6) and the downregulation of proteins related to sugar and glycogen processing (such as ACYP1, AK1A1, GALM and 6PGL) in FA Low-treated EHTs compared to CT High (Figure 3b, Table S4). When looking specifically at metabolic processes (Figure 3c), FA Low treatment promoted enhanced expression of proteins involved in the TCA cycle, FA β-oxidation, and mitochondrial energy homeostasis (from lower to higher contribution), while expression of proteins involved in glycolysis remained similar (Figure 3c, Table S5). Particularly, proteins related to FA β-oxidation, including ACADS, ACADV and CPT1A, and TCA cycle, including CISY and ODO1, were upregulated in EHTs treated with FA Low medium compared to control. This observed increase in the abundance of metabolic proteins is consistent with increased mitochondrial protein content, such as TOM70, CYC1, COX5B and ATP5PF (Table S5). Thus, both gene expression analysis and proteomic analysis further supported the increase in FA β-oxidation, accompanied by increased TCA cycle activity and a subsequent switch to a more oxidative energy metabolism.

Additionally, as increased OxPhos activity and cardiac maturation are associated with mitochondrial biogenesis ^44^, we examined the mitochondrial content and distribution using the mitochondrial marker TOM20 in EHTs cultured in FA Low medium compared to the control CT High medium. In both conditions, mitochondria were predominantly localized in the vicinity of sarcomeres, characteristic of more mature CMs ^45^ (Figure 3d). In line with the proteomics data, TOM20 intensity significantly increased in EHTs in FA Low medium, which suggested an increase in mitochondrial content (Figure 3e). Interestingly, although the gene expression of sarcomeric markers, such as myosin heavy chain 7 (*MYH7*) and troponin isoform 3 (*TNNI3*), was upregulated in FA-treated EHTs (Figure S4e), suggesting increased sarcomeric maturity, their corresponding protein levels remained unaltered (Figure S4f). In accordance, sarcomeric organization, observed by alpha-actinin (ACTN2), cardiac troponin T (TNNT2) and myosin light chain 2 (MLRV) staining, was comparable between conditions (Figure S4g-h). Similarly, Connexin 43 (CX43) localization and its punctate pattern were unchanged (Figure S4h), indicating that FA treatment did not induce overt morphological changes in tissue architecture while increasing mitochondrial abundance.

### 3.5. EHTs respond with improved functionality when cultured in FA Low medium and exhibit increased susceptibility to ischemic insults

A comprehensive functional evaluation was performed by tracking EHT movement over time with or without electrical pacing, and studying key parameters of cardiac kinetics: (1) force of contraction (peak), being the maximal force exerted by EHTs and representing mechanical output, (2) contraction velocity (CV, positive slope), and (3) relaxation velocity (RV, negative slope), which represent changes in the cardiomyocyte contraction and relaxation cycle, respectively.

In all tested cell lines, EHTs cultured in FA Low medium showed a decreased spontaneous beating frequency (Figures 4a-b and 4e-f), which is a hallmark of maturation. Proteomics data supported this by showing overall higher expression of proteins involved in calcium handling and electrophysiology (Figure 3c, Table S5), including proteins involved in regulating heart rate (Table S5). Because we observed differences in absolute contractile force between the hPSC lines used, derived from hESCs and hiPSCs, we next compared their responses to the FA Low condition. When electrically paced, the hESC line showed suboptimal contractile forces in CT High medium compared to the hiPSC line, which improved in FA Low medium (Figures 4c-d and S5a-b). In contrast, the hiPSC line already displayed high contractile force in CT High and maintained or further enhanced functional performance in FA Low, as reflected by higher or preserved contraction force as well as contraction and relaxation velocities (Figures 4g-h and S5c-d).

**Figure 4.**
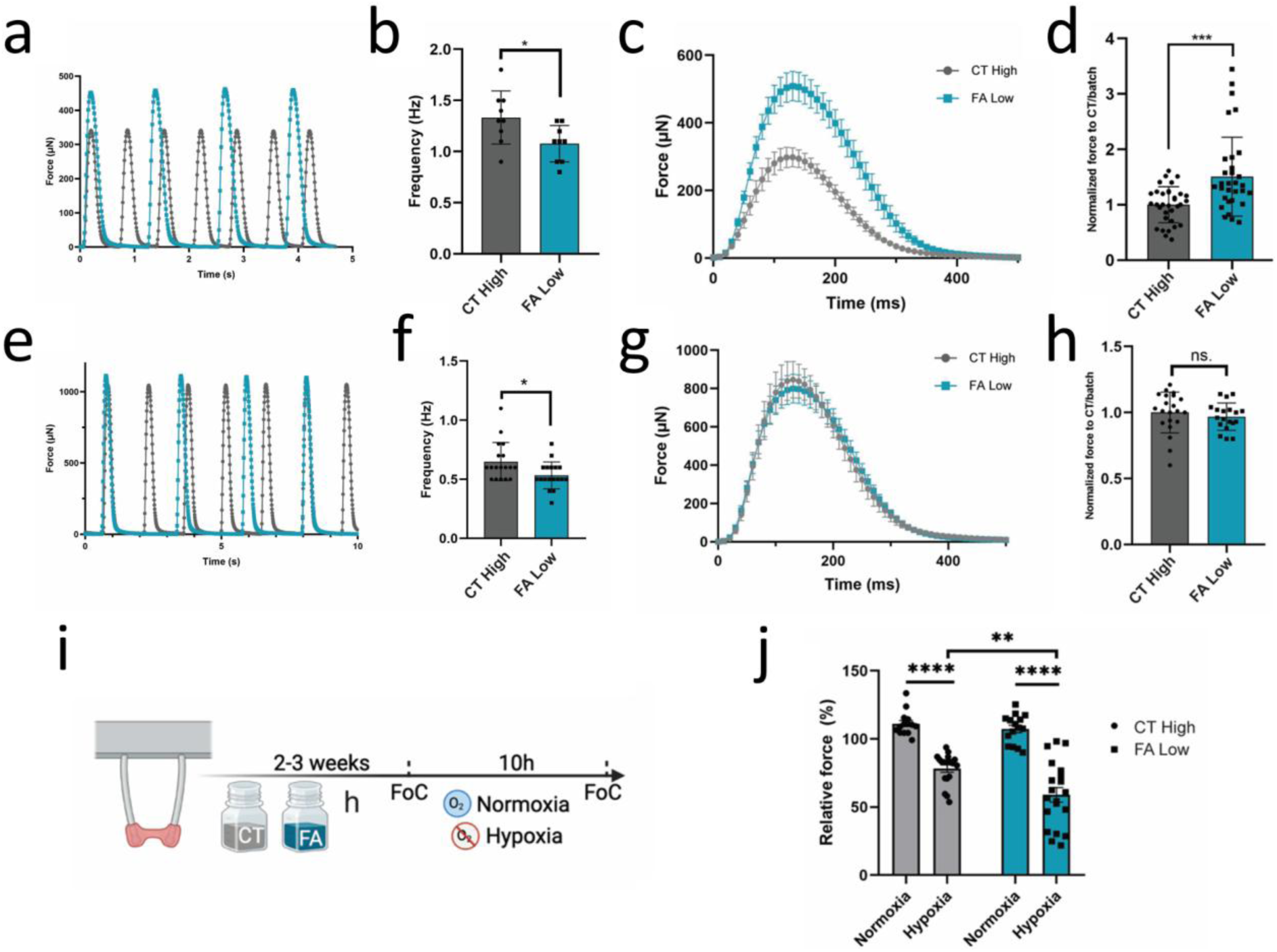
Metabolic maturation due to FA supplementation with physiological concentration of insulin promotes a functional improvement in EHTs, inducing increased sensitivity to ischemia insult. (a) Representative traces of contraction force over time in spontaneously recorded hESC-CM EHTs, cultured in CT High or FA Low medium. (b) Spontaneous beating frequency of EHTs cultured in CT High or FA Low medium (n = 9 tissues from 3 hESC-CM batches). (c) Representative averaged traces of the contraction peaks of paced hESC-CM EHTs, in CT High or FA Low medium (n = averaged contraction peaks of 3 tissues per condition). (d) Normalized contraction force to CT High per CM differentiation batch (n > 20 tissues from 4 hiPSC-CM batches). (e–h) Equivalent analyses in hiPSC-CM EHTs: (e) Spontaneous beating traces. (f) Spontaneous beating frequency (n = 18 tissues from 3 batches). (g) Representative paced contraction peaks (n = 3 tissues per condition). (h) Normalized contraction force (n = 19-21 tissues from 4 batches). Unpaired t-test with Welch’s correction (b,d,f,h). (i) Schematic representation of ischemia assay. Briefly, after 2-3 weeks of culture in CT High or FA Low medium, tissues were exposed to normoxic or hypoxic conditions for 10 hours, while maintained in their corresponding media. Before and after exposure, contraction force (FoC) was measured. (j) Relative contraction force normalized to baseline before normoxia/hypoxia exposure. Two-way ANOVA with Sidak post-hoc test (N = 14-20 tissues from 4 hiPSC-CM batches). *p<0.05; **p<0.01; *** p<0.0005; **** p<0.0001; ns = not significant.

To evaluate whether metabolic maturation in FA Low medium increases sensitivity to hypoxia, thereby enabling ischemia modeling, we assessed EHT functionality following hypoxia exposure. Before hypoxia exposure, baseline force of contraction measurements were recorded. After that, EHTs were exposed for 10 hours to hypoxia (0.1% O_2_) or normoxia (21% O_2_) (Figure 4i). Endpoint forces were normalized to each EHT’s baseline value. Under normoxia, EHTs in both media maintained contractile performance after 10 hours. In contrast, EHTs cultured in CT High medium exhibited a 22% reduction in force following hypoxia, likely reflecting their limited dependence on OxPhos for energy production (Figure 4j). In FA Low-treated EHTs, in which dependence on OxPhos was higher, hypoxia induced a more pronounced force reduction of 42%. These findings indicate that the metabolic shift induced by FA Low supplementation increases hypoxia sensitivity and, for the first time, enables ischemia modeling with a functional readout in hPSC-CM-derived EHTs.

### 3.6. EHTs demonstrated improved responses to positive inotropes and higher resistance to negative inotropes when cultured in FA Low medium

To assess drug response and predictability, we generated dose-response curves of FDA-approved inotropes and compared the potency of drugs on EHTs to clinically relevant plasma concentrations ^46–53^. Both CT High and FA Low-treated EHTs responded to inotropes, indicating high standard performance in both media (Figure 5).

**Figure 5.**
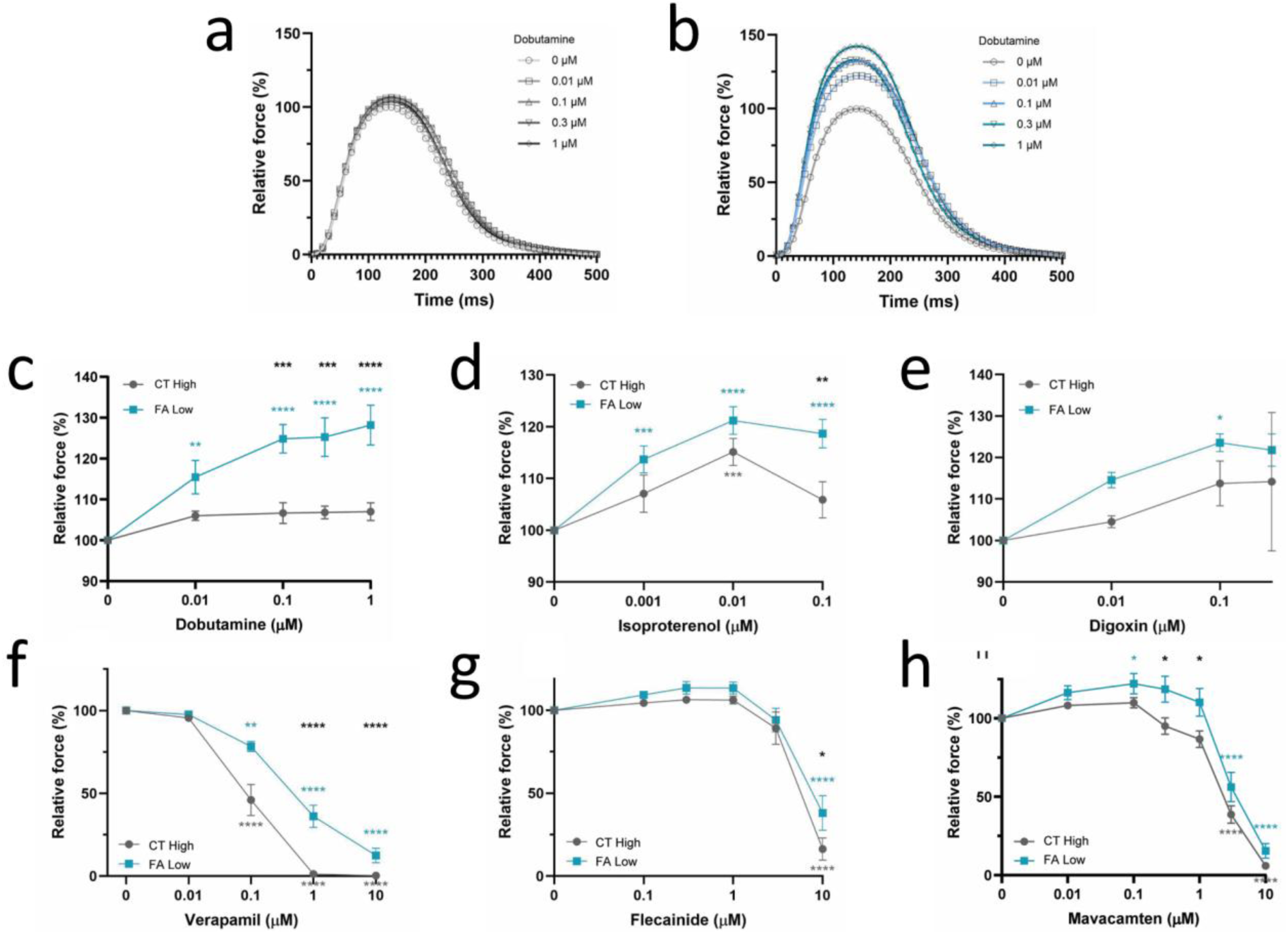
Improved drug predictability in metabolically matured EHTs. (a-b) Representative averaged traces of the contraction peaks of paced EHTs exposed to increasing concentrations of dobutamine in CT High (a) or FA Low (b) medium. (n = averaged contraction peaks of 3 EHTs per condition). (c-h) Relative contraction force response to increasing concentrations of dobutamine (a), isoproterenol (b), digoxin (c), verapamil (d), flecainide (e) and mavacamten (f) in EHTs treated with CT High or FA Low medium. Data are expressed as normalized percentages. Dose response curves were fitted using nonlinear regression (four-parameter logistic, variable slope), EC50 and IC50 values were calculated from the fitted curves. Two-way ANOVA with Sidak post-hoc test. (n = 9 tissues (a), 11 tissues (b), 9 tissues (c), 9 tissues (d), 12 tissues (e), 9 tissues (f), from 3-4 hiPSC-CM batches). Data plotted as means ± SEM. *p<0.05, **p<0.01, ***p<0.0005, **** p<0.0001. In (i-m), stars indicate significance within the same condition (light gray for CT High and light blue for FA Low), and between conditions (black).

Relative to the baseline measurement, EHTs cultured in CT High only showed a minor, non-significant effect to dobutamine, a β1-adrenergic receptor agonist, as shown by the relative force measured over time (Figure 5a), suggesting underdeveloped β1-adrenergic receptor expression or signaling. In contrast, EHTs cultured in FA Low medium exhibited a strong dose-dependent increase in peak contractile force in response to dobutamine, along with steeper contraction and relaxation slopes, indicating increased contraction and relaxation velocities (Figures 5b-c and S6a-c). From the dose-response in FA Low-treated EHTs, an EC50 of 10 nM was calculated (Figure S6a), which is an order of magnitude lower than reported mean efficacious plasma concentration in patients with congestive heart failure ^50^. EHTs cultured in both CT High and FA Low significantly improved cardiac kinetics in response to isoproterenol, a non-selective β-adrenergic receptor agonist (Figures 5d and S6d-i), with a stronger response in FA-treated EHTs and EC50 values of 1 nM and 0.9 nM, respectively (Figures 5d and S6g), matching reported plasma concentrations required for therapeutic effect ^52,53^. Greater improvements in force of contraction as well as contraction and relaxation velocities in FA Low-cultured EHTs upon dobutamine and isoproterenol treatment suggest an overall improved β-adrenergic response. In agreement, the expression of proteins involved in the β-adrenergic signaling cascade showed an overall enhancement in FA Low conditions (Figure S6j). This includes adenylyl cyclase ADCY6, an enzyme contributing to cyclic AMP (cAMP) synthesis ^54^, protein phosphatase 4 regulatory subunits (PP4P1 and PP4P2), which are involved in the regulation of protein phosphorylation states that control cardiac contractility and electrophysiology ^55^, and potential β-adrenergic signaling modulators such as the adhesion G protein-coupled receptor ^56^.

Upon stimulation with digoxin, a sodium/potassium-transporting ATPase inhibitor, FA Low-treated EHTs significantly improved cardiac kinetics (Figures 5e and S6k-m) with an EC50 of 15 nM (Figure S6k), corresponding to reported therapeutic ranges of serum concentrations in patients with heart failure ^51^. In contrast, CT High-treated EHTs did not show increased cardiac performance. Proteomic analysis revealed upregulation of digoxin targets AT1A2 and AT1A3 in FA Low-treated EHTs, consistent with their increased responsiveness (Tables S4 and S5). Verapamil, a voltage-gated L-type calcium channel blocker, significantly reduced cardiac kinetics in both CT High and FA Low conditions (Figures 5f and S6n-p). In comparison, FA Low-treated EHTs demonstrated increased resistance to verapamil, with an IC50 of 0.35 µM (Figure S6n), which falls within the reported plasma concentration range associated with therapeutic effects ^49^. This higher tolerance to Ca²⁺ channel blockade suggests more mature calcium handling and contractile regulation in FA Low-treated tissues. Flecainide, a sodium channel protein type V α subunit blocker, reduced the cardiac kinetics in both CT High and FA Low conditions with IC50 values within reported therapeutic plasma concentrations ^48^ (3.6 and 3.5 µM, respectively, Figure S6q). While the overall response was comparable, FA Low-treated EHTs exhibited slightly greater resistance (Figures 5g and S6q-s). These drug responses to digoxin, verapamil and flecainide suggest enhanced electrophysiology and calcium handling in FA Low-treated EHTs. Consistently, FA Low-treated EHTs exhibited faster calcium kinetics, showing a shorter time to peak in the hESC line with suboptimal performance in CT High medium and a similar trend in the high-performing hiPSC line (Figure S5e-h). Proteomic analysis further revealed overall higher expression of protein involved in calcium handling and electrophysiology (Figure 3c, Table S5). Specifically, NAC1, CHERP and CASQ1, three proteins involved in calcium homeostasis, were upregulated (Table S5) ^57–60^. Lastly, treatment with mavacamten, an inhibitor of the ATPase activity of cardiac myosin, reduced the cardiac kinetics in both CT High and FA Low conditions with IC50 values within reported target plasma concentration ranges to achieve therapeutic effects (1.9 and 2.4 µM, respectively, Figure S6t) ^46,47^. FA Low-treated EHTs displayed slightly greater resistance (Figures 5h and S6t-v), potentially due to increased actin-myosin functionality. Overall, EHTs cultured in FA Low showed improved pharmacological responses to FDA-approved inotropes, which aligned with the range of reported clinically-effective plasma concentrations, suggesting enhanced pharmacological maturation and improved predictability.

## 4. DISCUSSION

In this study, we demonstrate that supplementing 3D engineered heart tissues (EHTs) with fatty acids (FA) and physiological insulin levels promotes metabolic maturation, enhances pharmacological responsiveness and increases susceptibility to ischemic injury. By integrating proteomic and metabolomic profiling with functional characterization, we further demonstrate how metabolic remodeling influences contractile function, calcium handling, pharmacological sensitivity, and responses to hypoxic stress. Importantly, these findings extend beyond optimization of culture conditions and highlight the importance of metabolic maturation for improving the physiological relevance of engineered cardiac tissues in drug testing and disease modeling.

In agreement with other studies, FA supplementation improved hPSC-CM maturation ^6,61,62^. However, a key limitation in these studies is the widespread use of 2D cultures to assess metabolic parameters, such as respiratory capacity rates. While informative, metabolic readouts from 2D cultures may not accurately reflect the metabolic state of cells in a 3D environment due to their different energetic demands ^6^. In fact, Ulmer et al. reported that culture of hPSC-CMs in a 3D-EHT format alone is sufficient to enhance metabolic maturation, as indicated by an increased mitochondrial mass and number, upregulation of FA metabolizing enzymes and resemblance of mitochondrial biogenesis to that observed in the early postnatal heart ^10^. Furthermore, a recent study illustrated that EHTs exhibited a reduced action potential duration upon FA administration, while 2D monolayers remained unaltered ^6^. Therefore, in our study, we employed an integrated approach combining metabolomics, proteomics, and functional analyses to characterize metabolic remodeling directly in 3D-EHTs. This involved metabolomics profiling, respirometry and quantification of ATP levels in EHTs, revealing different metabolic dynamics across the four tested media conditions. Importantly, the integration of molecular profiling with functional measurements allowed us to investigate how changes in metabolic pathways are associated with cardiac performance, rather than relying exclusively on conventional maturation markers. We found that addition of FA enhanced overall TCA cycle activity. However, a pronounced switch to more oxidative metabolism only occurred when insulin concentration was reduced to physiological levels. These findings suggest that the effects of FA supplementation depend strongly on the broader metabolic environment and that insulin availability is an important determinant of metabolic adaptation in 3D cardiac tissues.

Insulin plays a key role in the regulation of multiple aspects of CM metabolism and function by mediating both glucose and long-chain FA metabolism. Since insulin concentration in the control medium far exceeds the physiological levels (17×10^3^ pmol/L in the ITS-X supplement vs 35-140 pmol/L in blood during fasting), it potentially represses FA uptake and OxPhos activity while promoting glucose uptake and metabolism ^63,64^ ultimately leading to increased lactate production. In contrast, physiological levels of insulin are permissive for the uptake and metabolism of both glucose and FA ^65,66^. This is consistent with the observed sustained glycolytic capacity and increased FA metabolism in the FA Low condition. This finding is also supported by a study in which complete removal of insulin and palmitate supplementation promoted higher contractility ^7^. Thus, the concentration of insulin should be carefully considered when optimizing the formulation of CM culture medium.

In this context, the choice of an appropriate control medium is crucial for enabling accurate and biologically relevant comparisons. For example, both our control media and FA-supplemented media contain physiological glucose levels (4.5 mM), as opposed to the high-glucose media commonly used to benchmark FA supplementation, which can induce cellular stress and impair CM maturation ^67,68^. To mitigate this and promote FA β-oxidation, glucose is often reduced or replaced with galactose exclusively in FA-based media, while control conditions remain unchanged ^5–9^. However, excluding glucose or substituting it with alternative sugars does not accurately recapitulate *in vivo* metabolism and complicates co-culture with glycolysis-dependent cell types such as fibroblasts and endothelial cells. Likewise, other metabolites such as lactate, carnitine ^69,70^, and taurine, are typically added only in FA-based media, even though they independently enhance hPSC-CM metabolism in 3D models ^71,72^. Moreover, Ca^2+^ levels critically regulate CM contractility, yet control media are often formulated at sub-physiological concentrations (below 1.6 mM) ^73^. Since EHT contraction force increases with rising Ca^2+^ levels until reaching a physiological plateau, maturation media containing higher Ca^2+^ concentrations cannot be directly compared to standard control conditions. Therefore, the higher percentage of FA supplement in our FA-based medium (3 times higher than the manufacturer’s recommended) may be attributed to the use of a highly specialized control medium, which we have previously demonstrated to enhance hPSC-CM maturation in EHTs. Importantly, this FA level brings the medium’s FA content closer to the physiological levels present in the adult human heart ^36^. Together, these considerations underscore the importance of controlling multiple medium components when interpreting metabolic maturation, particularly when comparing studies using different culture formulations.

Our findings also revealed that FA supplementation with physiological insulin levels led to a significant increase in contraction force and improved Ca^2+^ handling, with responses varying between cell lines according to differences in basic performance in CT High medium. This variability is consistent with previous studies, indicating that enhanced cardiac output of EHT does not uniformly occur under FA culture conditions ^6,7,62^, and that different hPSC lines may respond variably to metabolic cues ^74^. Nonetheless, FA Low treatment consistently reduced beating frequency in both hPSC lines tested, reflecting a more quiescent and mature cardiac phenotype ^75^. Beyond these functional maturation parameters, an important finding of our study is that metabolic culture conditions altered the pharmacological responses of EHTs. This provides a functional perspective on metabolic maturation, as changes in molecular and metabolic profiles become particularly relevant for preclinical applications when they are accompanied by altered responses to clinically relevant compounds. Drug testing in our system revealed that EHTs cultured in FA Low exhibited IC50 and EC50 values more closely aligned with human therapeutic plasma concentrations for several FDA-approved drugs, suggesting improved translatability. Although this comparison does not establish clinical predictivity, it supports the potential value of metabolic maturation for improving the pharmacological relevance of EHT-based assays.

Although sarcomeric protein expression remained unaltered, differences in inotropic responses observed with FA treatment could be attributable to an overall enrichment of calcium-handling and β-adrenergic related proteins, as well as improved mitochondrial biogenesis, in agreement with previous studies ^76,77^. The proteomic findings therefore provide a potential molecular basis for the observed changes in pharmacological responsiveness, linking metabolic remodeling to pathways involved in excitation-contraction coupling and adrenergic signaling. Some drugs, such as verapamil or isoproterenol, followed previously reported dynamics in hPSC-CM maturation, displaying decreased and increased sensitivity, respectively ^78^.

Notably, we also studied the response to mavacamten, a recently FDA-approved myosin ATPase inhibitor that has not yet been evaluated in the context of hPSC-CM maturation. We found that metabolically matured hPSC-CMs exhibited reduced sensitivity to mavacamten-induced contractile inhibition. Mavacamten binds allosterically to myosin, reducing actin affinity ^79^. This altered sensitivity may reflect differences in baseline contractile performance or the functional state of the contractile apparatus following metabolic maturation. However, further mechanistic studies are needed to establish the underlying explanation. Together, these pharmacological findings indicate that metabolic maturation can influence drug concentration-response relationships and should therefore be considered when developing and interpreting EHT-based drug screening assays.

Lastly, we demonstrated that our FA-based medium establishes a more physiologically relevant metabolic environment for evaluating ischemic injury. A particularly relevant finding is that metabolic maturation altered the response of EHTs to hypoxic stress, demonstrating that the metabolic state of engineered cardiac tissues can influence their susceptibility to disease-relevant injury. This is particularly important as ischemic injury in 3D cardiac models is commonly modeled by exposing hPSC-CM constructs to low pH media with high lactate levels under anoxic conditions for several hours, reflecting their predominant reliance on glycolysis for energy production ^36^. While this approach successfully recapitulates certain pathological features of ischemia, it does not fully capture the metabolic processes observed *in vivo*, where elevated lactate and acidosis are secondary consequences of ischemic injury rather than primary inducers of cellular damage ^80^. In contrast, our more metabolically mature, FA-dependent cardiac model showed increased susceptibility to hypoxic injury, consistent with an increased reliance on oxidative metabolism, thereby enhancing its relevance for studying ischemia-related molecular mechanisms. These findings highlight the importance of metabolic maturation in determining tissue responses to pathological stress. Future studies, such as modeling reperfusion injury following ischemia, or incorporating additional cardiac cell types to investigate tissue remodeling, will provide further insights into mechanisms of CM damage and support the identification of novel therapeutic approaches.

In conclusion, FA supplementation combined with physiological insulin levels induces a metabolic shift in EHTs that enhances their maturation state and aligns their metabolic profile more closely with that of the adult human heart, as supported by complementary proteomic, metabolomic, and functional analyses. Importantly, these metabolic changes also influence pharmacological responses and susceptibility to hypoxic injury. Collectively, these findings advance the translational value of EHTs, supporting more physiologically relevant models for drug testing and investigating cardiac disease mechanisms.

## AUTHORS’ CONTRIBUTION STATEMENT (CReditT)

C.C.F., C.F.K. and F.C. contributed equally. Conceptualization, C.C.F., R.P.; Methodology, C.C.F., C.F.K., F.C., H.J.C.T.W.; Investigation, C.C.F., C.F.K., F.C., E.E.N., A.v.R.; Writing, C.C.F., C.F.K., F.C; Reviewing and editing, J.M.R.A., E.E.N., J.V., H.W., V.S., M.C.R., R.P.; Funding acquisition, R.P., F.C., H.J.C.T.W., M.C.R., E.E.N.; Supervision, R.P., M.C.R., V.S.

## ACKNOWLEDGEMENTS AND FUNDING

R.P. acknowledges financial support from the European Research Council (ERC, Advanced Grant Heart2Beat, project number 101098372), ZonMw TOP grant (project number 91217061) and NWO through grant NWO/ENW-XL 2019.029. M.C.R. acknowledges financial support from the European Innovation Council (101099000-3DCardiacHTS, HORIZON-EIC-2022-TRANSITION-01). F.C. acknowledges financial support from Prinses Beatrix Spierfonds (“Op weg naar therapie” grant W.OR22-14) and Stichting Stofwisselkracht (“HoPE project”, grant 2022-03). Proteomics measurements were performed at the Radboud Technology Center for Mass Spectrometry supported by the Netherlands X-omics Initiative partially funded by NWO (project 184.034.019, H.J.C.T.W.). E.E.N. acknowledges financial support from the Danish Diabetes and Endocrine Academy, which is funded by the Novo Nordisk Foundation (grant number NNF22SA0079901). J.M.R.A. acknowledges the Netherlands Organ-on-Chip Initiative, an NWO Gravitation project (024.003.001) funded by the Ministry of Education, Culture and Science of the government of the Netherlands.

## SUPPLEMENTARY MATERIALS & DATA AVAILABILITY

Supplementary figures and tables, along with the omics datasets generated in this study have been deposited in the FigShare repository (FigShare.com), under the DOI:10.6084/m9.figshare.33801250 (link: https://doi.org/10.6084/m9.figshare.33801250), and are publicly available as of the publication date.

## ABBREVIATION LIST

*Cell types:*

2D: two-dimensional
3D: three-dimensional
3D-EHT(s): engineered heart tissue(s)
CMs: cardiomyocytes
FBs: human adult cardiac fibroblasts
hESCs: human embryonic stem cell
hPSCs: human induced pluripotent stem cells
hiPSC-CMs: human induced pluripotent stem cell-derived cardiomyocytes

*Culture media:*

BPEL: Bovine serum albumin/P Polyvinyl alcohol/Essential Lipids medium
CT High: CM medium containing supraphysiological insulin concentration
CT Low: CM medium containing physiological (lower) insulin concentration
E8: Essential 8 medium
FA High: CT High medium containing AlbuMax II Lipid-Rich BSA, Biotin, vitamin B12 and supraphysiological insulin concentration
FA Low: CT High medium containing AlbuMax II Lipid-Rich BSA, Biotin, vitamin B12 and physiological (lower) insulin concentration

*Materials & Methods:*

2-DG: 2-deoxyglucose
AEC: adenylate energy charge
cDNA: complementary deoxyribonucleic acid
Cys: carbamidomethyl
CV: contraction velocity
DAPI: 4′,6-diamidino-2-phenylindole
dda-PASEF: data dependent acquisition - Parallel Accumulation SErial Fragmentation
dia-PASEF: data independent acquisition - Parallel Accumulation SErial Fragmentation
DIA-NN: Data Independent Acquisition-Neural Network workflow
DMSO: dimethyl sulfoxide
EC50: half-maximal effective concentration
fps: frames per second
GFP: green fluorescent protein
IC50: half-maximal inhibitory concentration
ITS-X: Insulin-Transferrin-Selenium-Ethanolamine supplement KOH potassium hydroxide
KOSR: Knockout serum replacement
LFQ: label-free quantitation
MRM: multiple reaction monitoring mode
MS: mass spectrometry
Oligo: oligomycin
PFA: paraformaldehyde
RV: relaxation velocity
SEM: standard error of the mean
timsTOF: trapped ion mobility spectrometry-quadrupole time-of-flight mass spectrometer
UHPLC: ultra-high performance liquid chromatography

*Metabolites proteins and pathways:*

ACTN2: alpha-actinin
ADP: adenosine 5′-diphosphate
(c)AMP: (cyclic) adenosine 5′-monophosphate
ATP: adenosine 5′-triphosphate
CX43: connexin 43
FA: fatty acids
MLRV: myosin light chain 2
MYH7: myosin heavy chain 7
OxPhos: oxidative phosphorylation
PP4P1/PP4P2: protein phosphatase 4 regulatory subunits (1/2)
TCA cycle: tricarboxylic acid cycle
TNNI3: troponin isoform 3
TNNT2: troponin T2

